# Easybiotics: a GUI for 3D physical modelling of multi-species bacterial populations

**DOI:** 10.1101/412197

**Authors:** Jonathan Naylor, Harold Fellermann, Natalio Krasnogor

## Abstract

**Motivation:** 3D physical modelling is a powerful computational technique which allows for the simulation of complex systems such as consortia of mixed bacterial species. The complexities in physical modelling reside in the knowledge intensive model building process and the computational expense in calculating their numerical solutions. These models can offer insights into microbiology, both in understanding natural systems and as design tools for developing novel synthetic bacteria. Developing a robust synthetic system typically requires multiple iterations around the specify->design->build->test cycle to meet specifications. This process is laborious and expensive for both the computational and laboratory aspects, hence any improvement in any of the workflow steps would be welcomed. We have previously introduced Simbiotics (Naylor, 2017) a powerful and flexible platform for designing and analysing three-dimensional simulations of mixed species bacterial populations. Simbiotics requires programming experience to use which creates barriers to entry for use of the tool.

**Results:** In the spirit of enabling biologists who may not have programming skills to install and utilise Simbiotics, we present in this application note Easybiotics, a user-friendly graphical user interface for Simbiotics. Users may design, simulate and analyse models from within the GUI, with features such as live graph plotting and parameter sweeps. Easybiotics provides full access to all of Simbiotics simulation features, such as cell growth, motility and gene regulation.

**Availability:** Easybiotics and Simbiotics are free to use under the GPL3.0 license, and can be found at: https://bitbucket.org/simbiotics/simbiotics/wiki/Home

## 1 Introduction

Physical modelling can provide many insights into natural systems and aid in the design of synthetic cohorts. Full integration of computational tools in systems and synthetic biology workflows has not been fully realised despite the development of numerous modelling tools (Lardon, 2011; Gorochowski, 2012). We previously introduced Simbiotics (Naylor, 2017), a flexible platform for modelling bacterial populations. Despite the potential of the platform, barriers to use such as the requirement of programming knowledge prevent the platform being readily adopted by experimentalists and bioinformations. Here we present Easybiotics, a graphical user interface (GUI) allowing for the use of Simbiotics without programming experience. An easy to install process and minimal software dependencies allow for the use of Easybiotics in many contexts, enabling the development of 3D population models of distinct bacterial species. Here we exemplify use of the tool through developing a basic model of a growing bacterial population. Modelling tutorials and further elaboration on tool functionality and implementation can be found on the Simbiotics website, publications and corresponding user manual (Naylor, 2017). https://bitbucket.org/simbiotics/simbiotics/wiki/Home. Simbiotics was used by the Newcastle iGEM team who won a gold accolade at iGEM 2017 in Boston, it has been used for training of Newcastle’s MSc students on the Synthetic Biology and Systems Biology masters course and also applied to studying oral and other biofilms.

### 2 Easybiotics

Easybiotics is a graphical environment to design and run Simbiotics models. The tool gives access to the Simbiotics library of model features where they can be composed into a model specification and simulated with additional features such as live graph plotting and parameter sweeps in a few mouse clicks.

In Easybiotics models are defined by attaching library modules to a model specification. The model specification is visualised as an interactive tree structure (highlighted in red in Fig 1a), where the user can right click on specification nodes, such as the ’forces’ node (other examples highlighted in orange) to attach library modules, such as a frictional force (other examples highlighted in green). Modules are specific simulation features such as simulation domain size, physical forces and biological activity. Each module has a text ID which can be used to bind module functionality together. Module parameters can be set upon definition or by editing them in the property view (highlighted in blue). Data collection and analysis modules can also be attached to the model specification.

**Fig 1.**
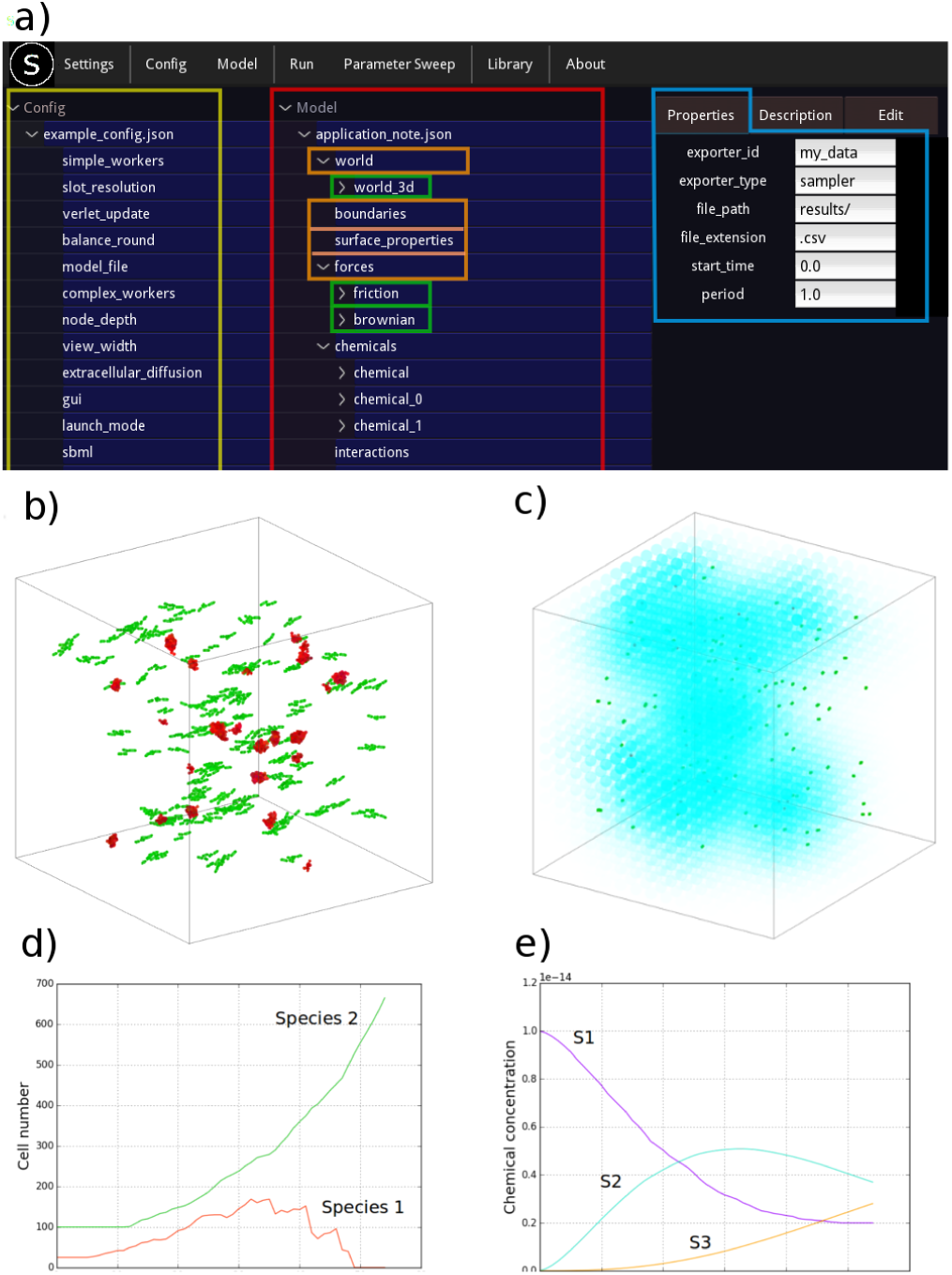
a) Easybiotics model development environment. The model specification tree (highlighted in red) is interactive and allows for adding and removing of library modules (modules highlighted in green). The configuration options for the Simbiotics simulator can be set (highlighted in yellow). The properties of the selected module (node on the specification tree) are displayed on the right panel, where a description of the module is also available. b) Snapshot of simulation at 30 minutes showing simulation domain with growing bacterial species 1 (red) andspecies 2 (green). c) Snapshot showing visualisation of chemical S2 concentration field at 10minutes. d) Cell numbers over time. e) Total chemical concentration over time

### 3 Developing and analysing biological models

An arbitrary number of cell species can be defined, where the metabolic behaviour of each species is defined separately. Each individual cell’s metabolic behaviour is decoupled from its species type, meaning each cell is distinct and can differentiate. Features for simulating intracellular dynamics include SBML models, boolean networks, sets of differential equations or stochastic simulation rules.

We exemplify the capacity of Easybiotics by building a hypothetical model and taking measurements of the simulation. The model consists of a coccus (spherical) and a bacillus (rod-shaped) species, both of which are planktonic suspended in a cubic volume of fluid, experiencing light mixing forces. To represent this we first define a 3D cubic simulation domain of length 100 *μ*m. A brownian motion and friction module are added to represent the mixing of cells in the fluid. We define a coccus and bacillus morphology, and the two distinct bacterial species. The species grow at different rates, and a constant nutrient source is assumed. Species 1’s metabolic pathway transforms chemical S1 into S2, and species 2’s metabolic pathway transforms S2 into S3. To represent the metabolic pathways we define an SBML model for each species. Both species have membrane transport systems to carry their respective reagents and products in and out of the cell. In addition, S3 in high concentrations is toxic to species 1, causing cell death. The system is induced by adding S1 to the extracellular space, which through diffusion and the cells metabolisms will eventually be transformed into S3 resulting in the death of species 1 cells. Initial conditions are defined describing the population of each cell species and amount of inducer chemical S1. Finally, data sampling is defined recording the total concentration of each chemical species and the cell number of bacterial species 1 and 2 every minute. Snapshots of the simulation and plots of the collected data can be seen in Figure 1.

### 4 Conclusion

The growth of the biotechnology industry is accelerating (Huggett, 2017). The application of CAD tools and automation to the industry is crucial for the mass-production of biodevices. Currently the design of distributed multicellular programs is a significant challenge due to the lack of methods developed. To partially alleviate these problems we have previously developed the Simbiotics modelling platform for the 3D modelling of bacterial populations. Here we have presented Easybiotics, a graphical user interface which removes barriers to entry for using the tool, making the integration of such computational modelling techniques into experimental workflows easier.

## References

Gorochowski, T.E. et al. (2012) BSim: AnAgent-Based Tool forModeling Bacterial Populations in Systems and Synthetic Biology. PLoS One, 7, e42790

Huggett et al. (2015) Research biotech patenting, Nature Biotechnology 34, 801–802

Lardon, L. A. et al. (2011) iDynoMiCS: next-generation individual-based modelling of biofilms Environ Microbiol., 13, 2416–2434,

Naylor, J. et al. (2017) Simbiotics: A Multiscale Integrative Platform for 3D Modeling of Bacterial Populations. ACS Synthetic Biology, 6, 1194–1210

